# Automated identification and quantification of stereotypical movements from video recordings of children with ASD

**DOI:** 10.1101/2024.03.02.582828

**Authors:** Tal Barami, Liora Manelis-Baram, Hadas Kaiser, Michal Ilan, Aviv Slobodkin, Ofri Hadashi, Dor Hadad, Danel Waissengreen, Tanya Nitzan, Idan Menashe, Analya Michaelovsky, Michal Begin, Ditza A. Zachor, Yair Sadaka, Judah Koler, Dikla Zagdon, Gal Meiri, Omri Azencot, Andrei Sharf, Ilan Dinstein

**Affiliations:** Department of Computer Science, Ben-Gurion University of the Negev, Beer Sheva, Israel; Azrieli National Centre for Autism and Neurodevelopment Research, Ben Gurion University of the Negev, Beer Sheva, Israel; Department of Psychology, Ben-Gurion University of the Negev, Beer Sheva, Israel; Department of Cognitive and Brain Sciences, Ben-Gurion University of the Negev, Beer Sheva, Israel; Department of Public Health, Ben-Gurion University of the Negev, Beer Sheva, Israel; Department of Computer Science, Bar-Ilan University, Ramat Gan, Israel; Pre-School Psychiatry Unit, Soroka University Medical Center, Beer Sheva, Israel; Zusman Child Development Center, Soroka University Medical Center, Beer Sheva, Israel; Child Development Center, Leumit Healthcare Services, Jerusalem, Israel; The Autism Center/ ALUT, Shamir (Assaf Harofeh) Medical Center, Be’er Ya’akov, Israel; Faculty of Medicine, Tel Aviv University, Tel Aviv, Israel; Neuro-Developmental Research Centre, Beer Sheva Mental Health Centre, Ministry of Health, Beer Sheva, Israel; Seymour Fox School of Education, The Hebrew University of Jerusalem, Jerusalem, Israel

## Abstract

**Importance:** Stereotypical motor movements (SMMs) are a form of restricted and repetitive behavior (RRB), which is a core symptom of Autism Spectrum Disorder (ASD). Current quantification of SMM severity is extremely limited, with studies relying on coarse and subjective caregiver reports or laborious manual annotation of short video recordings.

**Objective:** To demonstrate the utility of a new open-source AI algorithm that can analyze extensive video recordings of children and automatically identify segments with heterogeneous SMMs, thereby enabling their direct and objective quantification.

**Design, setting, and participants:** This retrospective cohort study analyzed video recordings from 319 behavioral assessments of 241 children with ASD, 1.4 to 8 years old, who participated in research at the Azrieli National Centre for Autism and Neurodevelopment Research in Israel. Behavioral assessments included cognitive, language, and autism diagnostic observation schedule, 2^nd^ edition (ADOS-2) assessments.

**Exposures:** Each assessment was recorded with 2-4 cameras, yielding 580 hours of video footage. We manually annotated 7,352 video segments containing heterogeneous SMMs performed by different children (21.14 hours of video).

**Main outcomes and measures:** We used a pose-estimation algorithm (OpenPose) to extract skeletal representations of all individuals in each video frame and trained an object-detection algorithm (YOLOv5) to identify and track the child in each movie. We then used the skeletal representation of the child to train an SMM recognition algorithm using a PoseConv3D model. We used data from 220 children for training and data from the remaining 21 children for testing.

**Results:** The algorithm accurately detected 92.53% of manually annotated SMMs in our test data with 66.82% precision. Overall number and duration of algorithm identified SMMs per child were highly correlated with manually annotated number and duration of SMMs (r=0.8 and r=0.88, p<0.001 respectively).

**Conclusion and relevance:** These findings demonstrate the ability of the algorithm to capture a highly diverse range of SMMs and quantify them with high accuracy, enabling objective and direct estimation of SMM severity in individual children with ASD. We openly share the “ASDPose” dataset and “ASDMotion” algorithm for further use by the research community.

**Key Points:** *Question:* Is it possible to train a deep learning algorithm to accurately identify and quantify stereotypical motor movements (SMMs) in video recordings of children with autism?

*Findings:* The ASDMotion algorithm was trained and tested with the largest video dataset of ASD children curated to date, comprised of 319 behavioral assessment recordings from 241 ASD children. The algorithm successfully detected 92.53% of manually identified SMMs with 66.82% precision, achieving highly accurate quantification of SMMs per child that were strongly correlated (r≥0.8) with quantification by manual annotation.

*Meaning:* This study demonstrates the utility of ASDMotion for objective and direct quantification of SMM severity in children with ASD, offering a new freely available, open-source algorithm and dataset that enable transformative basic and clinical ASD research.

## Introduction

Stereotypical motor movements (SMMs) are apparent in ∼50% of individuals with Autism Spectrum Disorders (ASD)^1^. They embody one form of restricted and repetitive behaviors (RRBs), which are a core symptom of ASD^2^. SMMs have been defined as repetitive, rhythmical, coordinated, seemingly purposeless movements^3–6^ that can be categorized into groups according to body topography^7^, complexity^8,9^, and/or function^10^. Common examples of SMMs include hand flapping, body rocking, jumping, and turning in circles. Although SMMs are not unique to ASD, they tend to be more prevalent in ASD than in other developmental disorders and typical development^3^.

SMMs may serve adaptive functions while causing maladaptive consequences. They are often described by individuals with ASD as a self-regulating coping mechanism for situations involving sensory overload, anxiety, or excitement^11–14^. However, frequent SMMs can disrupt learning, skill acquisition, and social communication, hindering integration into mainstream education or community settings^15,16^, and causing parental distress^17,18^. These divergent perspectives highlight a continuing debate about the clinical value and ethics of treatments for reducing SMMs ^10,19,20^.

Regardless of their function, SMMs must be reliably identified and quantified across different individuals and developmental stages to be studied. In previous studies, SMMs have mostly been measured using parent questionnaires such as the Repetitive Behaviors Scale-Revised (RBS-R)^21^, which includes a six-question stereotyped behavior subscale where parents rate the severity of specific SMM types. Although parent questionnaires provide an important perspective on SMMs, their accuracy and sensitivity is limited due to narrow scoring ranges and possible reporter bias^22.^

Other studies have measured SMMs in video recordings by manually annotating them^7,9,23,24^. While this approach allows direct quantification of SMMs, it is laborious, requires expertise, and results may be specific to the annotated video segment and its context. Studies using this approach have typically examined relatively short recordings of 10-20 minutes^7,23^. Additional studies applied machine-learning techniques to classify SMM types using Kinect^25,26^ (depth camera) recordings of SMMs performed in a lab setting or short video recordings (∼90-seconds) of SMMs captured by parents at home^27–29^. These studies, however, only attempted to distinguish between 3-4 predefined SMMs, rather than identify heterogeneous SMMs in lengthy video recordings of children with ASD.

Developing automated computer vision tools for identifying and quantifying SMMs in extensive video recordings would be transformative for the field, enabling direct, objective, scalable, high-throughput, low-effort quantification of this core ASD symptom. Achieving this task, however, requires overcoming two key challenges. First, most recordings contain more than one individual, making it necessary to identify and track the ASD individual so that only their movements are analyzed. Second, SMMs are highly heterogeneous, with ASD individuals displaying distinct and unique SMMs^30,31^. To accurately identify and quantify SMMs, it is, therefore, critical to train algorithms on large, well-annotated datasets that include both heterogeneous SMM exemplars and heterogeneous movements that are not SMMs.

To overcome these challenges, we curated the largest SMM dataset to date, composed of video recordings from 319 clinical assessments of 241 children. We extracted the skeletal representation (i.e., locations of 17 body joint positions) of all individuals visible in each frame, yielding a compact, sparse, and efficient representation of their body movements across frames. We then manually identified 7,352 video segments where the child exhibited an SMM, while labeling the skeleton of the child. This manually annotated library of SMMs, named ASDPose, was used to train and test the ASDMotion algorithm.

## Methods

### Participants

We analyzed video recordings from 241 children, ages 1.4-8-years-old, who were recruited between 2017 and 2021 at the Azrieli National Centre for Autism and Neurodevelopment Research (ANCAN), a collaboration between Ben-Gurion University of the Negev (BGU) and 8 clinical sites throughout Israel. The ANCAN autism database includes, among other measures, video, and audio recordings of clinical assessments ^32,33^. We selected children with a diagnosis of ASD according to DSM-5 criteria^2^ who completed an Autism Diagnostic Observation Schedule, 2^nd^ edition (ADOS-2) assessment^34^ where they scored two or more on ADOS-2 item D2 (repetitive hand and finger movements) and/or item D4/D5 (stereotypical behavior in ADOS-2 modules 1&2/Toddler’s respectively). The Helsinki committees at Soroka University Medical Center (SUMC), Shamir Medical Center, and Leumit Healthcare Services as well as the Internal Review Board of the Hebrew University approved this study. Parents of all participating children signed an informed consent form.

### Behavioral assessment recordings

We analyzed 883 video recordings from 319 behavioral assessments of 241 children. These included 226 ADOS-2 assessments (a semi-structured, standardized clinical assessment for identifying ASD symptoms^34^), 71 Preschool Language Scale, 4^th^ edition (PLS-4)^35^ assessments, and 22 developmental and cognitive assessments composed of 11 Mullen Scales of Early Learning^36^ assessments, and 11 Wechsler Preschool and Primary Intelligence, Third Edition^37^ assessments. Behavioral assessment rooms were equipped with 2-4 IP video cameras that recorded the assessments with a resolution of 1080x1920 pixels at 30 frames per second.

### Pose Estimation & Tracking

We used OpenPose^38^ to extract the locations of 17 skeletal joints in two-dimensional space for each person in each frame (typically, two or more individuals were in the room). Next, we applied a Spatial-Temporal Affinity Field tracking algorithm^39^ which assigned a unique number to each skeleton that remained the same across consecutive frames where the skeleton was visible and not partially occluded.

### Manual annotation of SMMs and the child’s skeleton

SMMs were manually identified and annotated using in-house developed software (Supplementary Figure 1). Annotators were undergraduate students trained by a clinician with >15 years of experience. They viewed the videos, labeled the start and end time of each SMM, classified its type (see Supplementary Table 1), and marked the skeleton ID of the child (a number corresponding to the skeleton’s color). This resulted in a list of 7,352 video segments containing SMMs that were, on average, 9.89 seconds long (median = 7.98 seconds).

### Computing hardware

Algorithm development, including all training and testing, was performed with two NVIDIA RTX3090 GPUs, a 32-core AMD Ryzen Threadripper PRO 3975WX CPU (3.5GHz), and 264GB of RAM.

### Child Detection

We trained the YOLOv5^40^ object detection algorithm, using a variant pre-trained on the COCO object dataset^41^, to identify the child in each video frame. To achieve this, we used 30,000 video frames from the 7,352 manually annotated SMM video segments described above where the child’s skeleton was manually labeled. Bounding boxes were constructed around the child’s skeleton and the skeletons of all other individuals (i.e., adults) in each frame. We then used 80% of the frames to train the algorithm and 20% for testing. The YOLOv5 algorithm achieved 95% precision and 92% recall for correctly identifying bounding boxes containing a child. We applied the algorithm to all frames of all movies and performed all further analyses only on child skeletons (Supplementary Figure 2).

### SMM identification algorithm (ASDMotion)

In addition to the 7,352 manually identified SMM video segments described above, we extracted 28,648 randomly selected video segments with an equivalent distribution of lengths that contained skeletal movements of children not identified as SMMs. We defined the data from the first 220 children (295 assessments) as our training dataset (6,597 segments with SMMs and 24,923 without) and the remaining 21 additional children (24 assessments) as our test dataset (755 segments with SMMs and 3724 without). There were no significant differences in the gender, age, or behavioral scores of children in the train and test datasets (Table 1).

**Table 1:** Descriptive statistics of sex, age, ADOS-2 scores, cognitive scores, and PLS-4 scores.

|  |  | Train set<br>N= 220 | Test set<br>N= 21 | Statistical Comparison |
| --- | --- | --- | --- | --- |
| Sex | Males | 172 (78%) | 15 (71.43%) | $\chi^2(1, N = 260) = 0.19, p = .65$ |
|  | Females | 48 (22%) | 6 (28.57%) |  |
|  | Mean (SD) |  |  |  |
| Age | | 3.97 (1.3) | 4.32 (1.39) | $t(258) = -1.19, p = .24$ |
| ADOS-2 | Total Calibrated Severity Score (CSS) | 7.07 (2.23) | 6.67 (1.96) | $t(258) = 0.8, p = .43$ |
| | Social Affect (SA) CSS | 6.52 (2.38) | 6.33 (2.17) | $t(258) = 0.35, p = .73$ |
| | Restricted Repetitive Behaviors (RRB) CSS | 8.01 (1.54) | 7.48 (1.54) | $t(258) = 1.51, p = .13$ |
| Cognitive | Developmental\Cognitive score | 76.49 (18.28)<br>(N=183) | 73.31 (13.83)<br>(N=16) | $t(214) = 0.68, p = .5$ |
| PLS- 4 | Total Standard Score | 68.18 (21.01)<br>(N=148) | 71 (14.33)<br>(N=9) | $t(174) = 0.4, p = .69$ |

We used a PoseConv3D model^42^ that was pre-trained on the Kinetics-400 dataset^43^ to create the ASDMotion algorithm. The PoseConv3D model uses a 3D heatmap of the skeletal movements as input to a 3D convolutional neural network (CNN), which was trained to identify SMMs. This architecture was specifically designed to identify spatial and temporal dependencies in human skeleton movements over time (i.e., compact representations of human body movements). The algorithm was trained with samples from the training dataset that were 200-frames long (i.e., 6.7 seconds). Training samples had a batch size of 64 input skeleton sequences (containing sequences with and without SMMs) and the training process utilized a stochastic gradient descent optimization for 100 epochs. ASDMotion output was a score between 0 and 1 corresponding to its confidence that the segment contained an SMM.

### Testing the algorithm

To test ASDMotion, we split each video recording into a sequence of overlapping epochs using a sliding window with a width of 200 frames and a step size of 30 frames (i.e., 170 frame overlap). ASDMotion scores for the overlapping epochs were transformed into a frame-wise time-course by selecting the maximum value across overlapping epochs (Figure1). To demonstrate the utility of the algorithm we classified frames with a score ≥0.85 (arbitrary threshold) as containing an SMM and concatenated contiguous neighboring SMM frames into an SMM movement. This process produced a list of automatically identified SMMs with their respective start and end times, enabling us to calculate the total number of SMMs, the number of SMMs per minute, and their median length per child (Supplementary Table 2).

**Figure 1:**
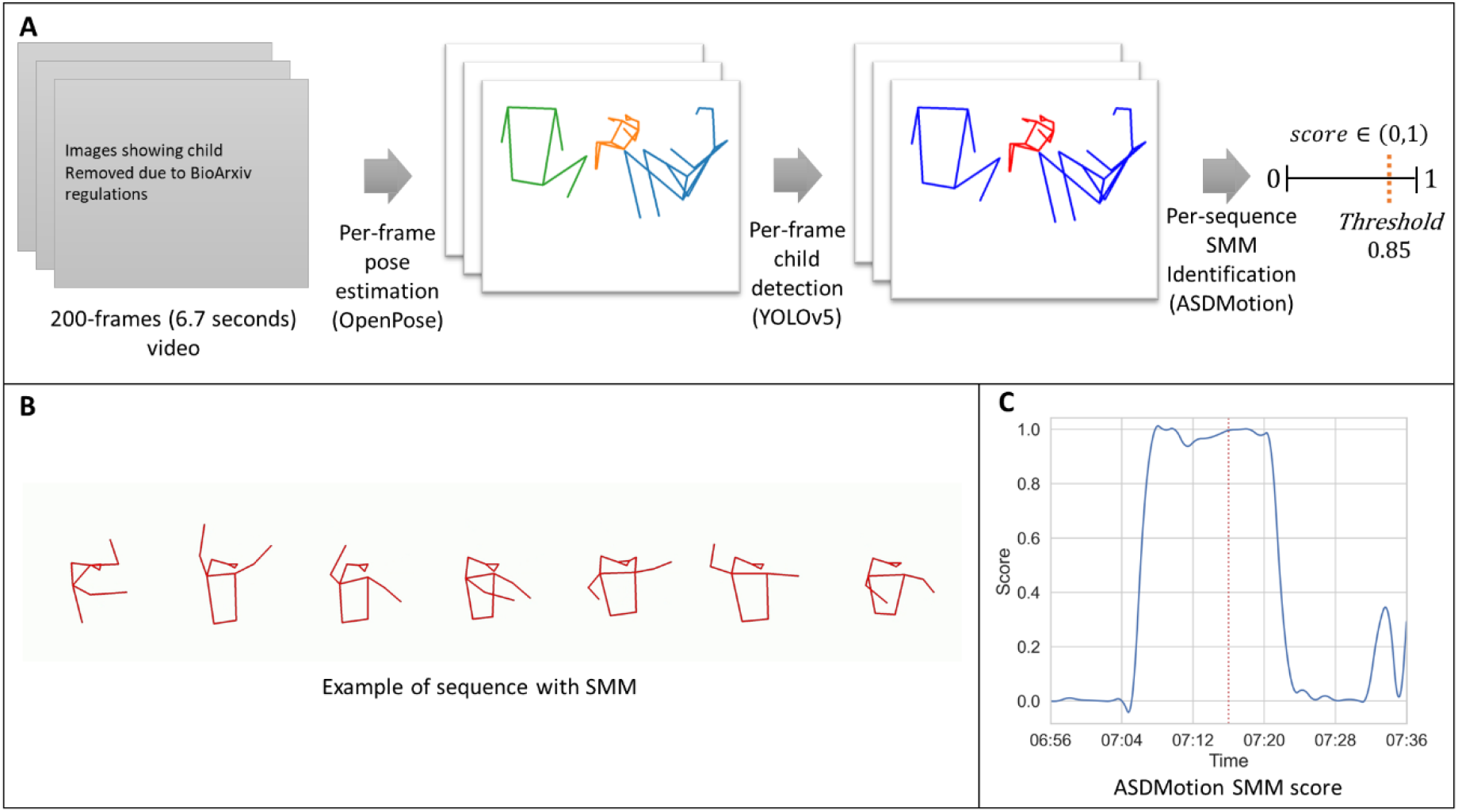
Schematic of algorithm pipeline and scoring. (A) Example of a 200-frame video segment that is processed with the pose estimation algorithm (OpenPose), then with the child detection algorithm (YOLOv5), and finally the child’s skeletal representation is analyzed by ASDMotion, yielding an SMM score between 0 and 1. (B) Example video and corresponding skeleton sequence where a child exhibits an SMM. (C) ASDMotion’s output for a 40-second video containing the SMM sequence in B. SMM scores are computed per 200-frame sliding window (6.7 seconds) with a 30 frame (1 second) step size, yielding a time-course of scores per video.

To assess accuracy, we calculated precision and recall per frame. Precision equals the number of true positive frames divided by the sum of true positive and false positive frames. Recall equals the number of true positive frames divided by the sum of true positive and false negative frames. Since SMMs are short, sparse, heterogeneous across children, and variable in their timing within each video, recall and precision values expected by chance are infinitesimal.

### Statistical analysis

All statistical analyses were performed using custom-written code in Python. Sex differences in precision and recall scores were evaluated using a Mann-Whitney test. Age and assessment score differences between the training and testing sets were assessed with independent sample t-tests. Pearson, Spearman, and concordance correlation coefficients were computed to assess the correspondence between the algorithm and the manual annotation. Finally, inter-rater reliability was estimated using percent agreement and Cohen’s Kappa. A statistical threshold of p>.05 was used throughout.

### Data sharing

The ASDMotion algorithm and ASDPose dataset are available for download at: https://github.com/Dinstein-Lab/ASDMotion

## Results

The number and duration of SMMs, as identified by manual annotation, varied greatly across children and behavioral assessments (Figure 2 and Supplementary Table 2). To adjust for different assessment lengths, we quantified the number of SMMs per minute of recording (Median: 0.12, Inter-quartile range = 0.06-0.22) and the percentage of time with SMMs per assessment (Median: 1.55%, Inter-quartile range = 0.64%-3.57%).

**Figure 2:**
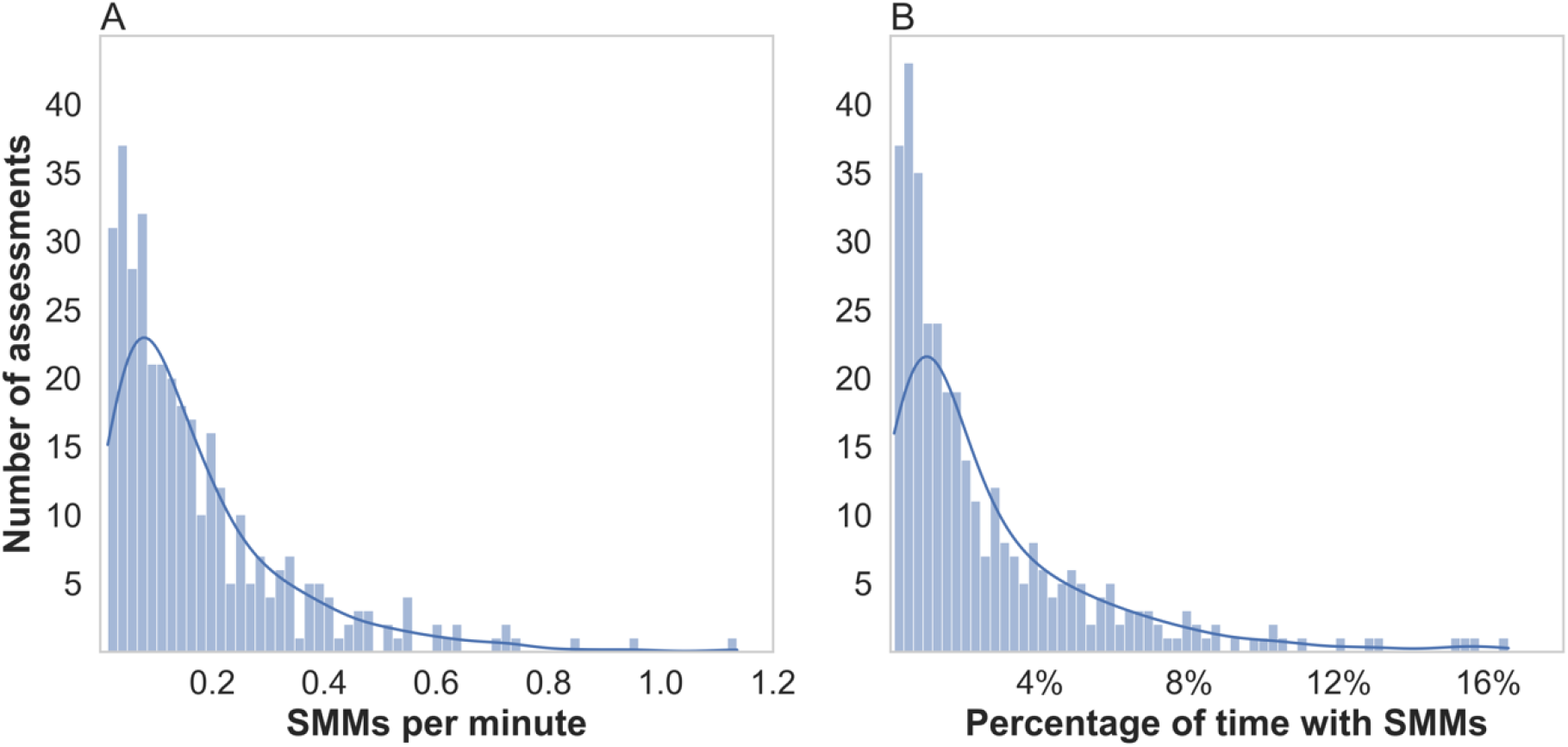
Different children with ASD exhibited different amounts of SMMs, as quantified by manual annotation. Histograms demonstrate the distribution of SMMs per assessment. (A) SMMs per minute. (B) Percentage of time with SMMs. Blue curve: probability density function (PDF) corresponding to each histogram.

### Initial accuracy of ASDMotion

After training ASDMotion with data from 295 assessments of 220 children (see Methods), we tested its accuracy with independent data from 24 assessments of 21 children who were not part of the training set (Figure 3A). We performed this analysis once with all children and again while separating males (n=15) and females (n=6). ASDMotion yielded a score between 0 and 1, representing the algorithm’s confidence about the presence of an SMM. We compared the algorithm’s scores with manual annotation per video frame while thresholding SMM classification at different score values. Frame-wise precision increased, and recall decreased when changing the classification threshold from a score of 0.5 to 0.9. For example, using a threshold of 0.85 yielded precision and recall of 36.64% and 87.72% for the entire group, 35.56% and 87.77% for males, and 39.93% and 87.59% for females, respectively. The high recall values suggested that ASDMotion accurately identified most manually annotated SMMs, but the low precision values indicated a high number of false positives. There were no significant differences in the accuracy of ASDMotion across males and females at the 0.85 threshold (precision: *U* = 23, *p* = 0.13, recall: *U* = 48, *p* = 0.54).

**Figure 3:**
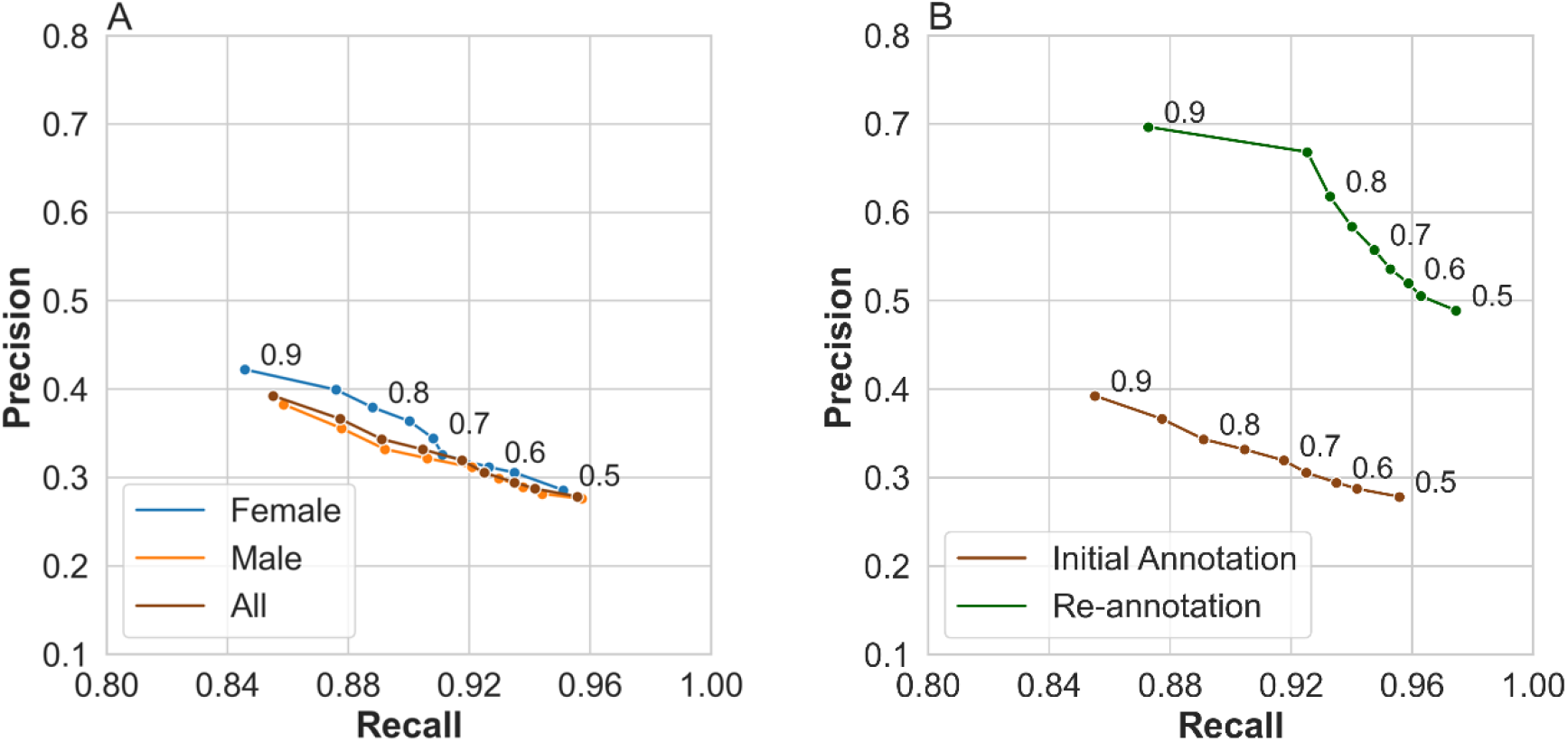
Precision and recall values demonstrate the accuracy of ASDMotion in identifying SMMs on individual video frames in the test data comprised of 24 assessments from 21 children. Each point represents precision and recall values when selecting a specific SMM classification threshold (i.e., algorithm scores from 0.5 to 0.9) (**A**) First analysis using the initially annotated 755 SMM and 3724 non-SMM segments in the test data. This analysis was performed once with all children (brown) and again with male (orange) or female (blue) children only. (**B**) Second analysis using only the 1456 video segments (728 SMM and 728 non-SMM) that were re-annotated by the two independent annotators who exhibited high test-retest reliability. Precision and recall values are presented for these video segments when using the initial manual labels (brown) and again when using the correct re-annotated labels (green).

### Re-annotation and inter-rater reliability of the test data

The low precision of the algorithm suggested that there may have been video segments with SMMs that were missed by the initial manual annotation. Concurrently, we also wanted to assess inter-rater reliability, which was not examined in the initial annotation process. To achieve both, we performed a second round of manual annotation with two independent annotators. We extracted 1,456 video segments from the 24 assessments in the test set that contained an equal number of true positives (hits), false positives (false alarms), true negatives (correct rejections), and false negatives (misses) in terms of the match between ASDMotion labeling and the initial manual annotation. The two annotators, blind to the initial annotation and the algorithm’s labeling, manually re-annotated these segments independently (i.e., labeled each segment as containing an SMM or not). There was 90% agreement between the two annotators, corresponding to excellent inter-rater reliability (𝜅 = 0.76).

Video segments identified as containing an SMM by either annotator were re-labeled as SMMs. The re-annotation process revealed that 51% of the video segments initially designated as false positives were actually true positives (i.e., contained an SMM that was missed in the initial annotation) and 9.8% of the segments initially designated as false negatives were actually true negatives (i.e., did not contain an SMM). We believe that the high percentage of missed SMMs demonstrates the difficulty of manually annotating SMMs within long video recordings of behavioral assessments. Testing the algorithm’s accuracy with the re-annotated 1,456 video segments revealed a true precision and recall of 66.82% and 92.53%, respectively, when using a threshold of 0.85 (Figure 3B).

### Accuracy of SMM quantification per assessment/child

While frame-wise precision-recall curves are important for determining the accuracy of ASDMotion in computer science terms, this assessment is overly conservative for basic and clinical autism research purposes where one is interested in quantifying the overall amount or rate of SMMs that a child exhibits. A more relevant accuracy test for such purposes would be to compare the number and duration of SMMs exhibited by each child as quantified by the SMM algorithm versus manual annotation (Figure 4). ASDMotion derived measures were strongly and significantly correlated with manually annotated measures for both the number of SMMs per assessment (r(22)=0.80, ρ(22)=0.80, ccc=0.70, p-value<0.001) and their percentage of time per assessment (r(22)=0.88, ρ(22)=0.87, ccc=0.73, p-value<0.001) when using Pearson, Spearman, or concordance correlation coefficients respectively. Note that this analysis utilized the re-annotated data described above.

**Figure 4:**
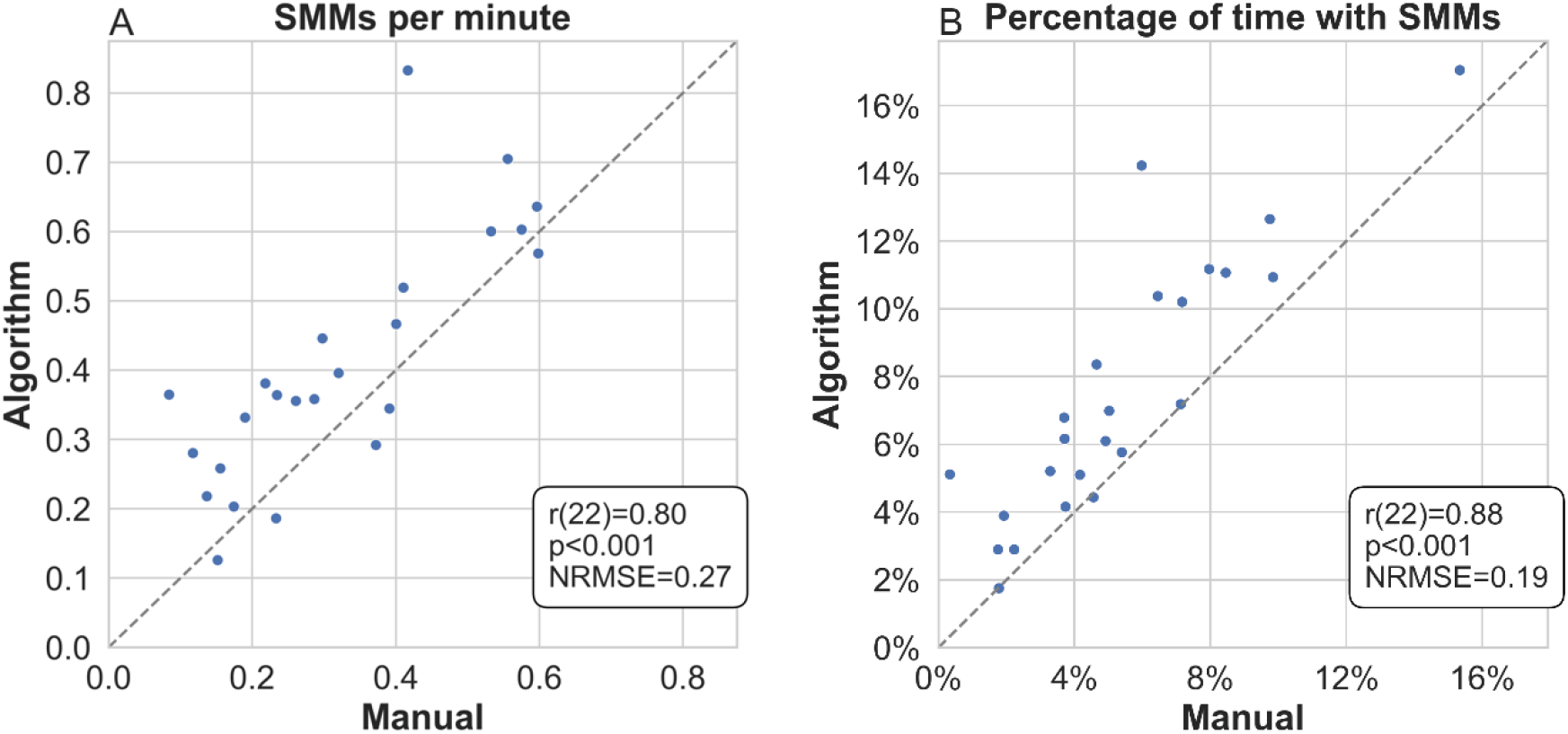
Accuracy of the algorithm in quantifying SMMs per assessment. Scatter plots compare the number of SMMs per minute (left) and percentage of time with SMMs (right) as quantified by manual annotation versus the algorithm. Pearson correlation coefficients and their statistical significance are noted in each panel.

## Discussion

ASDMotion was able to identify/recall >90% of the video frames that were manually labeled as containing SMMs. Initially, ASDMotion appeared to yield a high number of false positives (i.e., low precision, Figure 3A), but comprehensive re-annotation of 1,456 video segments by two independent annotators, who exhibited 90% agreement and high inter-rater reliability, revealed that 51% of segments initially considered false positives were in fact true positive, thereby increasing precision from 37% to 67%. This demonstrated that the annotators missed many SMMs during their initial examination of the long, ∼40-minute, assessment videos. These manual labeling errors were corrected only during re-inspection of the video segments marked by ASDMotion. This highlights the advantage of using an automated algorithm for detecting sparse events within long video recordings that are difficult and boring to manually annotate. Note that the re-annotation process was performed in a fair manner, by including an equal number of true/false positive and true/false negative segments in the 1,456 re-examined segments.

Most importantly, strong, significant correlations were evident between algorithm-derived and manually annotated SMM rates and durations per assessment (i.e., the number of SMMs per minute and the proportion of time with SMMs, Figure 4). These correlations demonstrate the value of ASDMotion for quantifying SMM severity per child. We believe ASDMotion has the potential to replace manual annotation techniques previously applied to short recordings in small samples^7,9,23,24^, thereby enabling large scale studies on a variety of topics, such as characterizing the development of SMMs in children with ASD and identifying their behavioural^43^, physiological^14^, and neural triggers.

### Previous SMM related algorithms and ASDMotion

The ASDMotion algorithm is novel and distinct from previous SMM classification algorithms. The algorithm can scan long video recordings of behavioral assessments and detect segments that contain a wide variety of heterogeneous SMMs (see Supplementary Table 1). In contrast, previously published algorithms were trained with short ∼90 second home-videos to distinguish between 3-5 specific types of SMMs such as hand flapping, head banging, and toe walking^27–29^. Additional studies trained algorithms to perform the same task while using Kinect depth camera recordings^26,28^, which also allow extraction of skeletal representations^44^. However, these algorithms only distinguished between predefined SMMs and are not able to identify heterogeneous, sparse SMMs within extensive real-life videos.

### Previous SMM related datasets and ASDPose

Previous examples of SMM data sets include curated short home-videos of SMMs that were recorded by parents of ASD children and posted online. The original Self-Stimulatory Behavior Dataset^45^ includes a selection of 75 videos (∼90 seconds long) containing examples of arm flapping, head banging, or spinning SMMs. The Expanded Stereotype Behavior Dataset^29^ includes a different selection of 141 videos (∼20 seconds long) containing spinning, arm flapping, hand action, and head banging SMMs. Neither dataset contains any clinical or demographic information about the recorded children, who may or may not have a formal diagnosis of ASD, and most recordings contain only the exhibited SMM.

In contrast, the ASDPose dataset contains the skeletal representation of ASD children in extensive recordings from 319 behavioral assessments (∼40 minutes long) of 241 children who have been thoroughly clinically characterized. The dataset includes demographic information as well as ADOS-2 scores for all the children and cognitive and PLS-4 scores for most. However, to maintain privacy, ASDPose does not include the raw video of these assessments and is limited to the child’s skeletal representation. The dataset is released along with the manual annotation of 7,352 SMM segments and details about our selection of training and testing data for transparency and to enable reproducibility. We expect that the availability of such extensive data from SMM and non-SMM behaviors will accelerate motion tracking research and development of additional algorithms for analyzing child behavior regardless of ASD.

### Limitations

The current version of the SMM algorithm has several limitations. First, it was trained only with video recordings from clinical behavioral assessments and may, therefore, produce less accurate results with videos recorded in other clinical or experimental contexts. Second, the current algorithm was not trained to classify between different types of SMMs. Third, the algorithm was trained only with videos of 1.4-8-year-old children and may, therefore, produce less accurate results with recordings of older individuals with ASD. All these limitations will be addressed in future versions of the algorithm as we and others train it on additional data.

## Conclusions

The ASDMotion algorithm and ASDPose dataset offer an exciting, revolutionary way of studying SMMs in ASD and other disorders when individuals exhibit SMMs. This novel digital phenotyping technique offers opportunities for studying the underlying physiological mechanisms that generate SMMs, for identifying the behavioral and neural events that trigger SMMs, and for determining the effectiveness of clinical interventions that aim to reduce SMMs. Future versions of the algorithm will extend its utility, robustness, and applicability even further.

