## Supplemental Information for "Automated identification and quantification of stereotypical movements from video recordings of children with ASD"

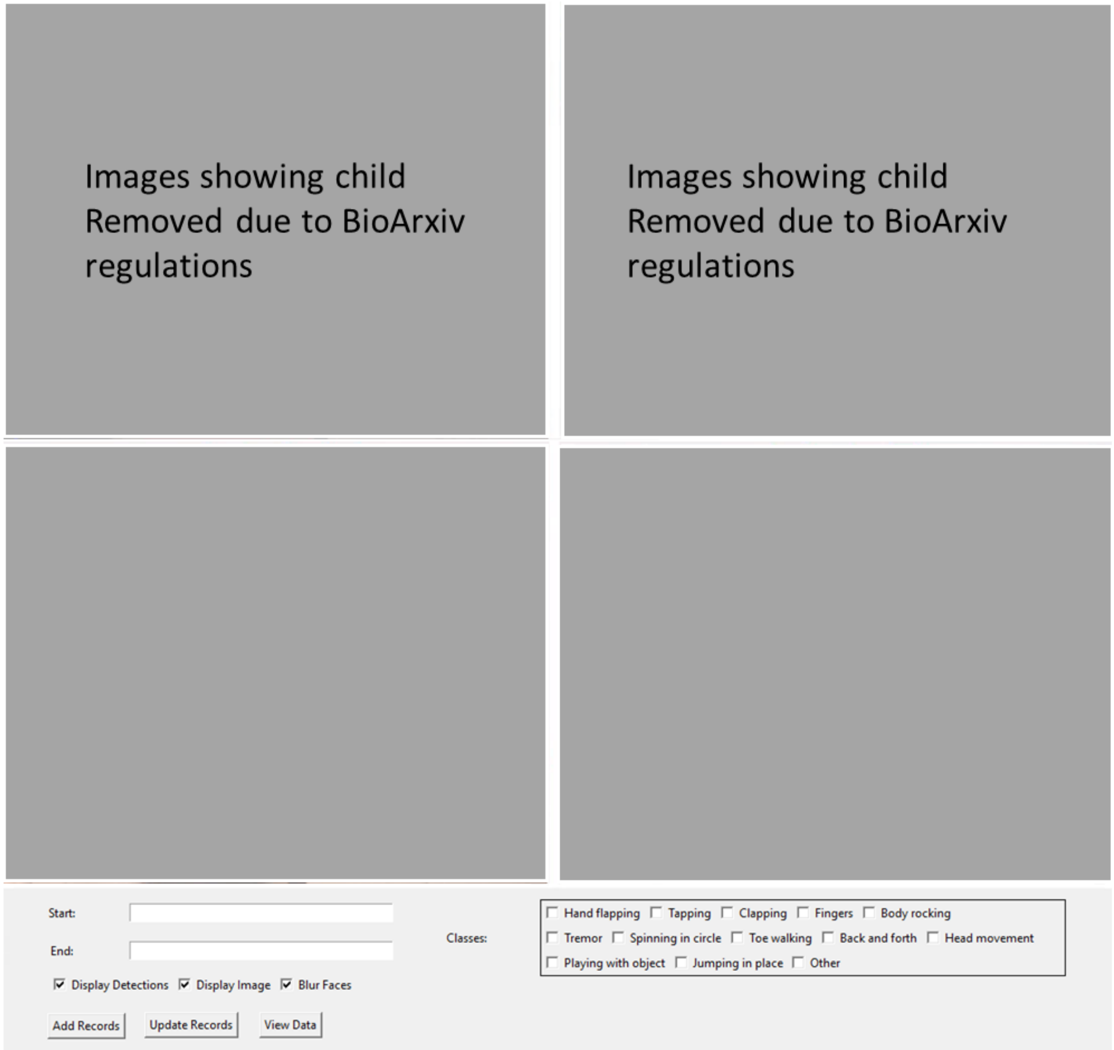


**Supplementary Figure 1:** Manual annotation was performed with in-house developed software that enabled annotators to quickly mark the start and end times of SMMs, the child's skeleton number, and the relevant cameras (i.e., views) where the child was visible.

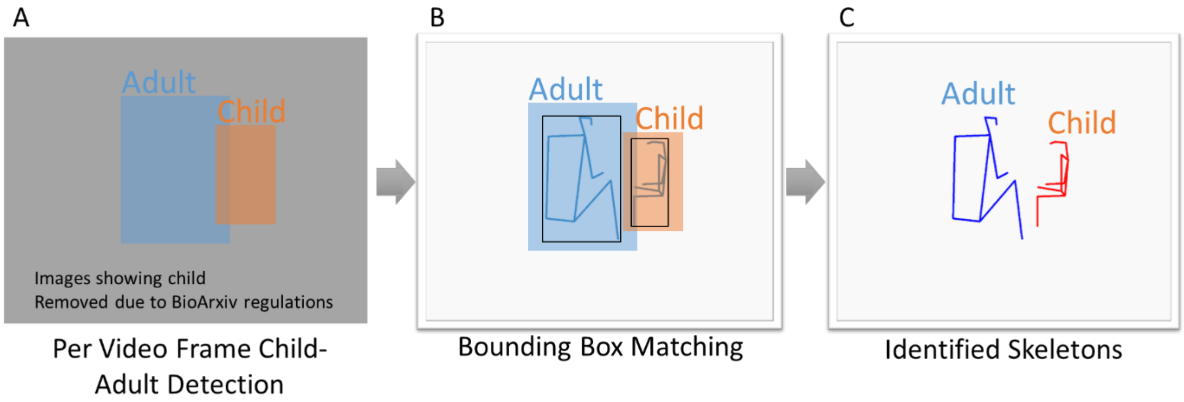


**Supplementary Figure 2:** Using YOLOv5 to identify child and adult skeletons. (A) The YOLOv5 object detection model was trained to identify all children and adults in each video frame. The algorithm marks the location of each with a rectangular bounding box. (B) Overlay of the YOLOv5 bounding boxes and the extracted OpenPose skeletons for the same video frame. Black rectangles are bounding boxes of each skeleton. (C) YOLOv5 child/adult labels are transferred to the OpenPose skeleton. Identity of each skeleton is determined by maximizing the intersection-over-union of the YOLOv5 and OpenPose bounding boxes. YOLOv5 bounding boxes that do not match any OpenPose skeleton are discarded, and skeletons with no matching YOLOv5 bounding box are labeled as Adult.

**Supplement Table 1:** List of SMM categories that were manually annotated and included in the current study. Categories were defined according to the physical characteristics of the movements.

| ***Category*** | **Description** |
|---|---|
| *Clapping* | Striking palms of hands against one another |
| *Hand flapping* | Flapping hands by moving the wrists or elbows up and down or side to side in the air |
| *Finger flicking* | Moving fingers in the air in a repetitive manner |
| *Tapping* | Repeatedly tapping a surface with the hand or fingers |
| *Spinning* | Spinning in circles while standing |
| *Pacing* | Walking back and forth along the same path |
| *Jumping* | Jumping or hopping repeatedly in the same spot or around the room |
| *Toe walking* | Walking on the toes instead of flat feet |
| *Body rocking* | Swaying the body back and forth or side to side while sitting or standing |
| *Tremor* | Rhythmic muscle contractions and relaxations yielding repetitive twitching movements of one or more body parts |
| *Playing with an object* | Using a toy in an unusual and repetitive manner. For example, repeatedly licking a ball or repeatedly spinning the wheel of a toy car. |
| *Head movement* | Rhythmic head movement from side-to-side or up-and-down |
| *Other* | A singular movement that was unique for a child or observed rarely and did not fit one of the categories above. For example, repeatedly rotating a yamaka, playing with hair in a peculiar way, repeatedly touching a point on the wall, etc.. |

***Supplement Table* 2:** Descriptive statistics of manually annotated video recordings.

| | **Median** | **IQR** |
|---|---|---|
| Assessment videos length (minutes) | 35.9 | 30.6-48.21 |
| Average duration of an SMM (minutes) | 0.13 | 0.08-0.19 |
| Percentage of video time with SMMs | 1.4% | 0.63%-3.21% |
| Total number of SMMs per video | 4 | 2-7 |
| SMM frequency per minute | 0.1 | 0.05-0.2 |
